# Multi-omics and Pharmacological Characterization of Patient-derived Glioma Cell Lines

**DOI:** 10.1101/2023.02.20.529198

**Authors:** Min Wu, Ran Yuan, Nan Ji, Ting Lu, Tingting Wang, Junxia Zhang, Mengyuan Li, Penghui Cao, Jiarui Zhao, Guanzhang Li, Jianyu Li, Yu Li, Yujie Tang, Zhengliang Gao, Xiuxing Wang, Anhua Wu, Wen Cheng, Ming Ge, Gang Cui, Yongping You, Wei Zhang, Qianghu Wang, Jian Chen

## Abstract

Glioblastoma (GBM) is the most common brain tumor and is currently incurable. Primary GBM cultures are widely used tools for screening potentially therapeutic drugs; however, there is a lack of genomic and pharmacological characterization of these primary GBM cultures. Here, we collected 52 patient-derived glioma cell (PDGC) lines and characterized them through whole-genome sequencing (WGS), RNA-seq, and drug response screening. We identified three molecular subtypes among PDGCs: mesenchymal (MES), proneural (PN), and oxidative phosphorylation (OXPHOS). Upon profiling the responses of PDGCs to 214 drugs, we found that the PN subtype PDGCs were sensitive to tyrosine kinase inhibitors, whereas the OXPHOS subtype PDGCs were sensitive to histone deacetylase inhibitors, oxidative phosphorylation inhibitors, and HMG-CoA reductase inhibitors. PN and OXPHOS subtype PDGCs stably formed tumors *in vivo* upon intracranial transplantation into immunodeficient mice, while most MES subtype PDGCs were incapable of tumorigenesis *in vivo*. In addition, profiling and follow-up investigations showed that the serum-free culture system used for PDGCs enriched and propagated rare *MYC/MYCN*- amplified glioma cells. Our study provides a resource for understanding primary glioma cell cultures and aiding clinical translation.

**Significance:** Our study provides a resource for patient-derived glioma cell lines (PDGCs) on transcriptome, genome, drug response, and tumorigenic abilities. PDGCs are categorized into PN, MES, and OXPHOS subtypes, with MES-subtype PDGCs incapable of tumorigenesis *in vivo*. Notably, the serum-free culture system for PDGCs enriches glioma cells with *MYC/MYCN* amplification.

## Introduction

Glioblastoma (GBM) is the most common malignant tumor of the central nervous system (CNS) ^1, 2^. The clinical translation and treatment options of GBM remain a huge challenge ^3^. Currently, the standard of care therapy for GBM is maximal surgical resection followed by temozolomide chemotherapy and adjuvant radiotherapy ^1, 2, 4^. The median survival of GBM is only about 15 months, and this has not improved significantly in the past few decades ^1, 5^.

Human cancer cell lines are the most commonly used models for mechanistic studies and drug screening in oncology research. Patient-derived glioma cells (PDGCs) cultured in serum-free neural stem cell medium have been widely used in glioma studies ^6, 7^. Compared to traditional glioma cell lines cultured in a serum-containing medium, PDGCs more faithfully recapture the mutational spectrums and gene expression patterns of parental tumors ^8, 9^. Several large-scale studies, including the Cancer Cell Line Encyclopedia (CCLE) ^10,11,12^, Genomics of Drug Sensitivity in Cancer (GDSC) ^13^, and Cancer Therapeutics Response Portal (CTRP) ^14^, have thoroughly characterized the genetic features and pharmacological responses of traditional glioma cell lines, while data depicting the genomic and pharmacological landscape for PDGCs is relatively scarce.

To address this challenge, we here integrated whole-genome sequencing (WGS), RNA-seq, and drug response data to characterize more than 50 PDGCs derived from glioma patients. Our data support the classification of the examined PDGCs into three molecular subtypes: mesenchymal (MES), proneural (PN), and oxidative phosphorylation (OXPHOS). Drug response profiles for 214 FDA-approved small molecules revealed that the PN subtype PDGCs were sensitive to tyrosine kinase inhibitors, while the OXPHOS subtype PDGCs were sensitive to histone deacetylase inhibitors, oxidative phosphorylation inhibitors, and HMG-CoA reductase inhibitors. Notably, PDGCs cultured in the serum-free medium did preserve the genomic alterations, subtype identity, and informative intra-tumor heterogeneity of parental tissues, however, the serum-free culture system enriched and propagated rare *MYC/MYCN*-amplified glioma cells. In addition, MES subtype GBM patients have a worse prognosis compared to the PN and OXPHOS subtypes, but we noted that the majority of cultured PDGCs with MES subtype failed to establish tumors upon intracranial transplantation into immunodeficient mice. Our study provides an integrated resource to support glioma-related studies.

## Results

### Establishment and multi-omics profiling of 52 patient-derived glioma cell lines cultured in serum-free medium

We collected 52 patient-derived glioma cell (PDGC) lines cultured in serum-free medium, which were derived from one diffuse intrinsic pontine glioma (DIPG), one low-grade glioma (grade II), and 50 GBMs. We performed RNA-seq on 52 PDGCs, whole-exome sequencing (WES) on 10 PDGCs, and whole-genome sequencing (WGS) on 51 PDGCs, as well as RNA-seq on 12 matched GBM tumor tissues of PDGCs (**Fig. 1A and Table S1**).

**Figure 1.**
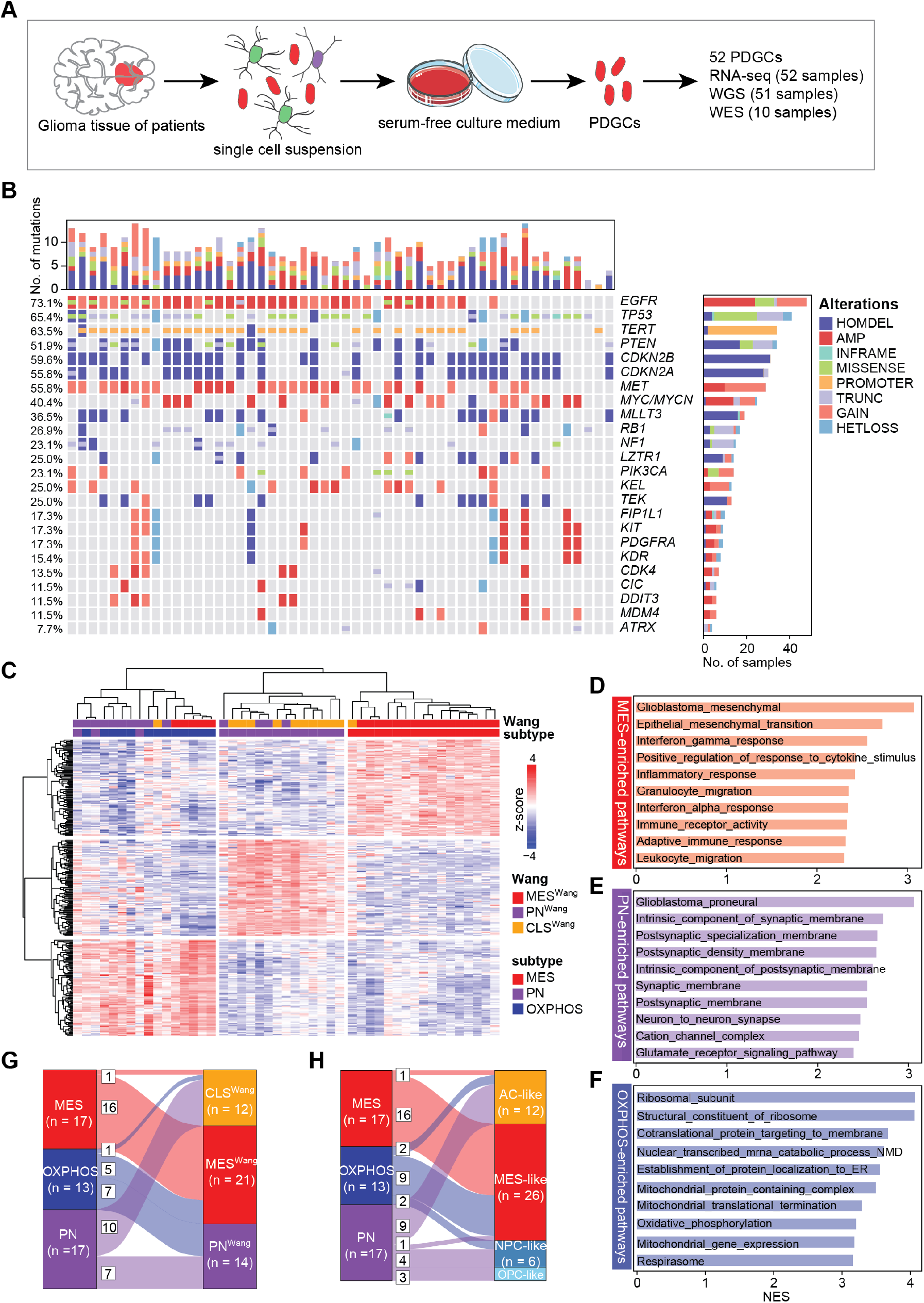
Definition of transcriptional subtypes for PDGCs, see also Figure S1. **(A)** Schematic diagram showing the generation of PDGCs. Glioma tissues from patients were dissected and digested into single-cell suspension, then cultured in a serum-free medium to obtain PDGCs. A total of 52 PDGCs were collected or generated in this study, of which 52 PDGCs were subjected to RNA-seq, 10 PDGCs were subjected to WES, and 51 PDGCs were subjected to WGS. **(B)** Oncoprint showing the genomic alteration of 52 PDGCs. HOMDEL: homozygous deletion, AMP: high-level amplification, INFRAME: inframe insertion and deletion, MISSENSE: missense mutation, PROMOTER: mutation located in promoters, TRUNC: truncation mutation, GAIN: low-level gain, HETLOSS: heterozygous deletion. **(C)** Heatmap showing the expression levels of subtype-specific signatures. Gene expression values were normalized by z-score. Rows represent genes, and columns represent PDGCs. **(D-F)** Barplots displaying the enriched pathways by MES (**D**), PN (**E**), and OXPHOS (**F**) subtype. Bars were colored by the subtype identity. ER: endoplasmic reticulum, NMD: nonsense-mediated decay. **(G-H)** Sankey plots showing the subtype assignment (n = 47) change flow. The left column is for our defined subtypes and the right column is for subtypes defined by Wang *et al*. (2017) (**G**) or subtypes defined by Neftel *et al*. (2019) (**H**).

Previous study ^8^ reported that glioma cells cultured in a serum-free medium more faithfully recapitulate the transcriptional patterns of parental tissues than cells cultured in a serum-containing medium. As a confirmation, we used genes that were highly expressed in GBM tissues and genes that were highly expressed in normal brain tissues to define a GBM tumor index (i.e., GBM score / normal score) (see **Methods**). A higher GBM tumor index represented a higher degree of similarity to GBM tissues. By integrating transcriptome profiles from The Cancer Genome Atlas (TCGA) and CCLE, we found that PDGCs cultured in a serum-free medium showed a higher GBM tumor index than glioma cell lines cultured by a serum-containing medium from CCLE (median GBM tumor index, TCGA-GBM: 3.06, PDGCs: 2.19, CCLE-GBM: 1.09, CCLE-glioma: 1.08) (**Fig. S1A**). It has been reported that primary glioma cell lines under a serum-containing medium have difficulty in maintaining the amplification of *EGFR* ^15^. Our WGS analysis revealed that PDGCs retained amplification of the *EGFR* and other common driver events of GBM ^9, 16^, including for example amplification of the *PDGFRA* genes, deletion of the *PTEN* and *CDKN2A* genes, and mutations in the *TP53* and *NF1* genes (**Fig. 1B**).

### Definition of transcriptional subtypes of PDGCs

Depending on their transcriptomes, GBMs are classified into three subtypes: classical (CLS^Wang^), proneural (PN^Wang^), and mesenchymal (MES^Wang^) at the bulk tumor level ^17^. At the single-cell level, GBM tumor cells have been classified into four cellular states: oligodendrocyte progenitor-like (OPC-like), neural progenitor-like (NPC-like), astrocyte-like (AC-like), and mesenchymal-like (MES-like) ^18^. Seeking to categorize PDGCs subtypes, we applied non-negative matrix factorization (NMF) to RNA-seq data for the 52 PDGCs and found that the four-cluster option achieved the highest cophenetic score (**Table S2**). We identified differentially expressed genes and constructed a 100-gene signature for each NMF-defined cluster (**Table S3 and Fig. 1C**).

Gene set enrichment analysis (GSEA) revealed that PDGCs in cluster 1 (n = 17) were enriched for epithelial-mesenchymal transition (EMT) and several immune-associated pathways (**Fig. 1D and Fig. S1B**), so we designated this cluster as the mesenchymal (MES) subtype. PDGCs in cluster 2 (n = 17) were enriched for neuron developmental-associated pathways (**Fig. 1E and Fig. S1B**), and we designated this cluster as the proneural (PN) subtype. PDGCs in cluster 4 (n = 13) were enriched for mitochondria-associated functions and the oxidative phosphorylation pathway (**Fig. 1F and Fig. S1B**), and we designated this cluster as the oxidative phosphorylation (OXPHOS) subtype. As cluster 3 only contained five cell lines and we did not find gene signatures that were significantly differentially expressed compared to other clusters, we mainly focused on the other three clusters. To check whether our subtypes were applicable to PDGCs from published studies, we examined the expression pattern of our signatures (**Table S3**) in the dataset from Mack *et al*. (2019) ^19^ and found our signatures also differentially expressed in these PDGCs (n = 44) (**Fig. S1C**).

To examine the relationship between our subtypes and the two existing subtyping approaches ^17, 20^, we assigned subtype identities of 52 PDGCs using all three subtyping approaches (**Table S3**). Our MES subtype exhibited enrichment in MES^Wang^ and MES-like state (16/17, 94.1%), indicating the mesenchymal signature was stable across the three subtyping approaches (**Fig. 1G, H**). Our PN subtype exhibited enrichment in PN^Wang^ (7/17, 41.2%) and CLS^Wang^ (10/17, 58.8%) and correlated with progenitor cell states (NPC- and OPC-like) (7/17, 41.2%). Our OXPHOS subtype mainly overlapped with PN^Wang^ (7/13, 53.9%) and MES^Wang^ (5/13, 38.5%) and correlated with MES-like cellular state (9/13, 69.2%) (**Fig. 1G, H**).

### PDGCs retain the subtype identity and heterogeneity of the parental tissues

To assess the extent to which PDGCs could retain the subtype identity of parental tissues, we used the RNA-seq data for 12 GBM tissues and their matched PDGCs to evaluate changes upon culture. We found that 66.7% (8/12) of PDGCs retained the subtype of their parental tissues upon culture; the four exceptions were one MES tissue transformed into OXPHOS PDGC, one OXPHOS tissue transformed into PN PDGC, and two PN tissues transformed into MES PDGCs (**Fig. 2A and Table S4**). A similar trend was evident when we examined another published dataset ^19^, wherein 70% (7/10) of PDGCs retained the subtype of their parent tissues, with one MES and two PN tissues transformed into OXPHOS PDGCs (**Fig. 2B and Table S4**). These two lines of evidence both supported that most of PDGCs retained the subtype identity of their parental tissues.

**Figure 2.**
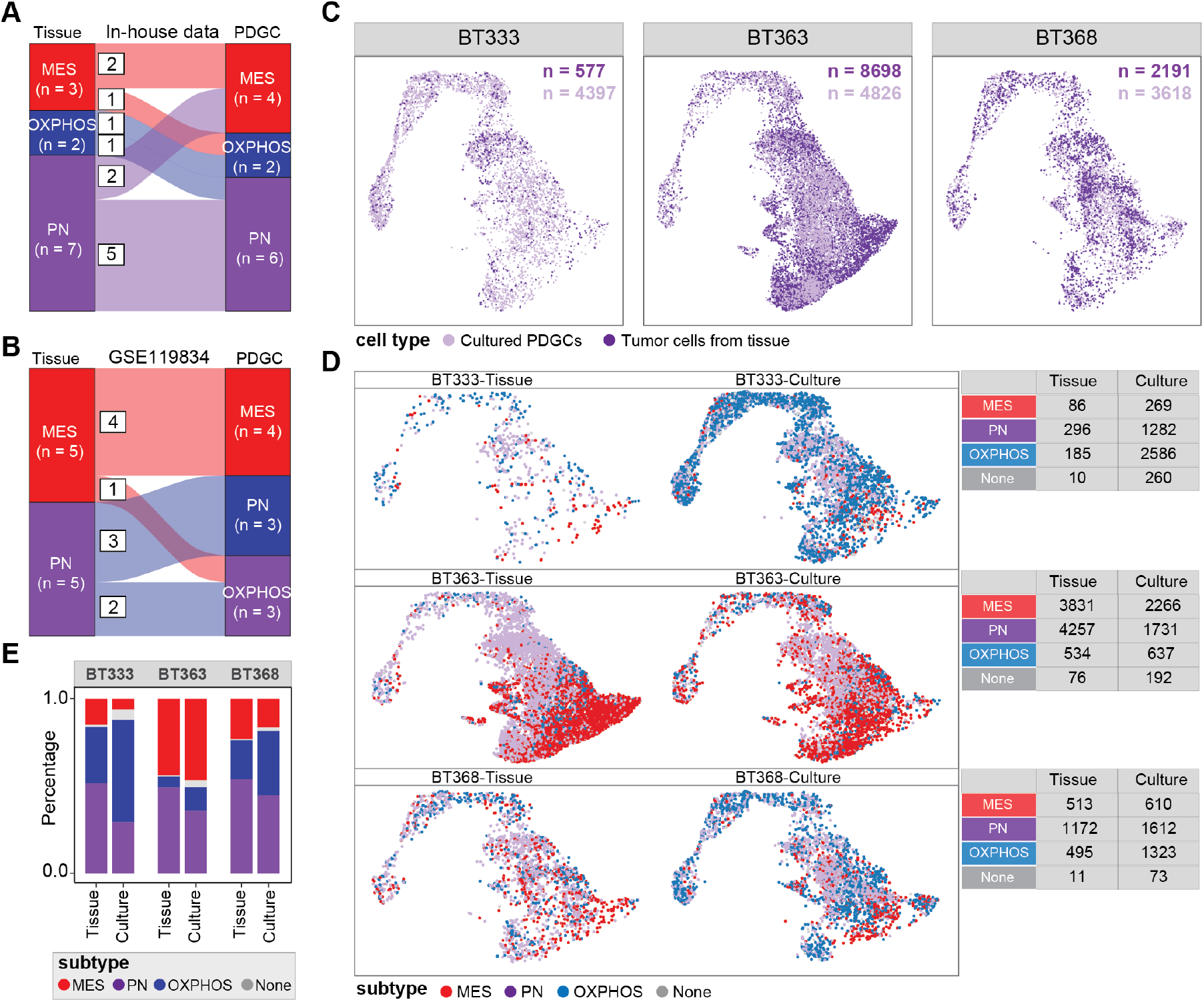
Subtype of cultured PDGCs and their original tissues, see also Figure S2. **(A-B)** Sankey plots showing the subtype assignment change between cultured PDGCs and their original tissues from in-house data (**A**) and previously published data (**B**). **(C)** Uniform manifold approximation and projection (UMAP) visualization of GBM tumor cells from tissue and cultured PDGCs, colored by cell type. **(D)** UMAP visualization of GBM tumor cells from tissues and cultured PDGCs, colored by subtype assignment. Tables next to the UMAP plot showing a brief summary of the cell number of each subtype. **(E)** The percentage of each subtype in GBM tumor cells from tissues and cultured PDGCs.

GBM is well known for its heterogeneity, and previous single-cell RNA-seq (scRNA-seq) analyses have demonstrated the co-occurrence of multiple tumor cell subtypes within a single tumor tissue ^20, 21^. To evaluate the degree to which cultured PDGCs can retain the heterogeneity of the parental tumor tissues, we analyzed scRNA-seq data from the Couturier *et al*. (2020) study, which assessed both cultured PDGCs and their matched tumor tissues ^22^. We initially filtered out non-malignant cells, and then re-clustered the tumor cells for the heterogeneity analysis (**Fig. S2**). A UMAP embedding indicated that the tumor cells isolated from tissues clustered with cultured PDGCs, supporting some extent of transcriptional similarity between the two sample types (**Fig. 2C**). And similar to the tumor tissues, cultured PDGCs derived from a single tumor tissue contained different subtypes of cells (**Fig. 2D**).

We subsequently compared the cell composition of different subtypes between cultured PDGCs and their parental tissues. For patient BT333, the percentage of cells with MES and PN subtypes decreased, while the percentage of cells with OXPHOS subtype increased in the cultured PDGCs (tumor tissue vs. cultures PDGCs: MES, 14.9% vs. 6.1%; PN, 51.3% vs. 29.1%; OXPHOS, 32.1% vs. 58.8%). Similar trends that MES and PN subtype tumor cells decreased and OXPHOS subtype tumor cells increased were observed for PDGCs from patient BT363 (MES, 44.0% vs. 47.0%; PN, 48.9% vs. 35.9%; OXPHOS, 6.1% vs. 13.2%) and BT368 (Tumor tissue vs. cultures PDGCs: MES, 23.4% vs. 16.9%; PN, 53.5% vs. 44.6%; OXPHOS, 22.6% vs. 36.6%) (**Fig. 2E**). These findings support that while the cultured PDGCs do preserve the multi-subtype tumor heterogeneity of the GBM parental tissues, their subtype composition does change slightly.

### Clinical, genomic, and tumorigenic comparisons of PDGCs with different subtypes

To examine the relationship between our classification system and the subtyping approach defined by Wang *et al*. ^17^ in GBM tissues, we classified GBM samples from the TCGA project using both subtyping strategies. Similar to what we have observed in PDGCs, our MES subtype exhibited enrichment in MES^Wang^ subtype (116/155, 74.8%), our PN subtype exhibited enrichment in PN^Wang^ (108/231, 46.8%), and CLS^Wang^ (116/231, 50.2%), while our OXPHOS subtype distributed in all the three subtypes defined by Wang *et al*. (**Fig. S3A, S3B**). Additionally, survival analysis showed that GBM patients of the MES subtype had significantly shorter overall survival (OS) compared to patients of the other two subtypes (median OS: 12.6 months for the MES subtype, 14.7 months for the PN subtype, and 15.1 months for the OXPHOS subtype, log-rank test, *p* = 0.0099) (**Fig. 3A**).

**Figure 3.**
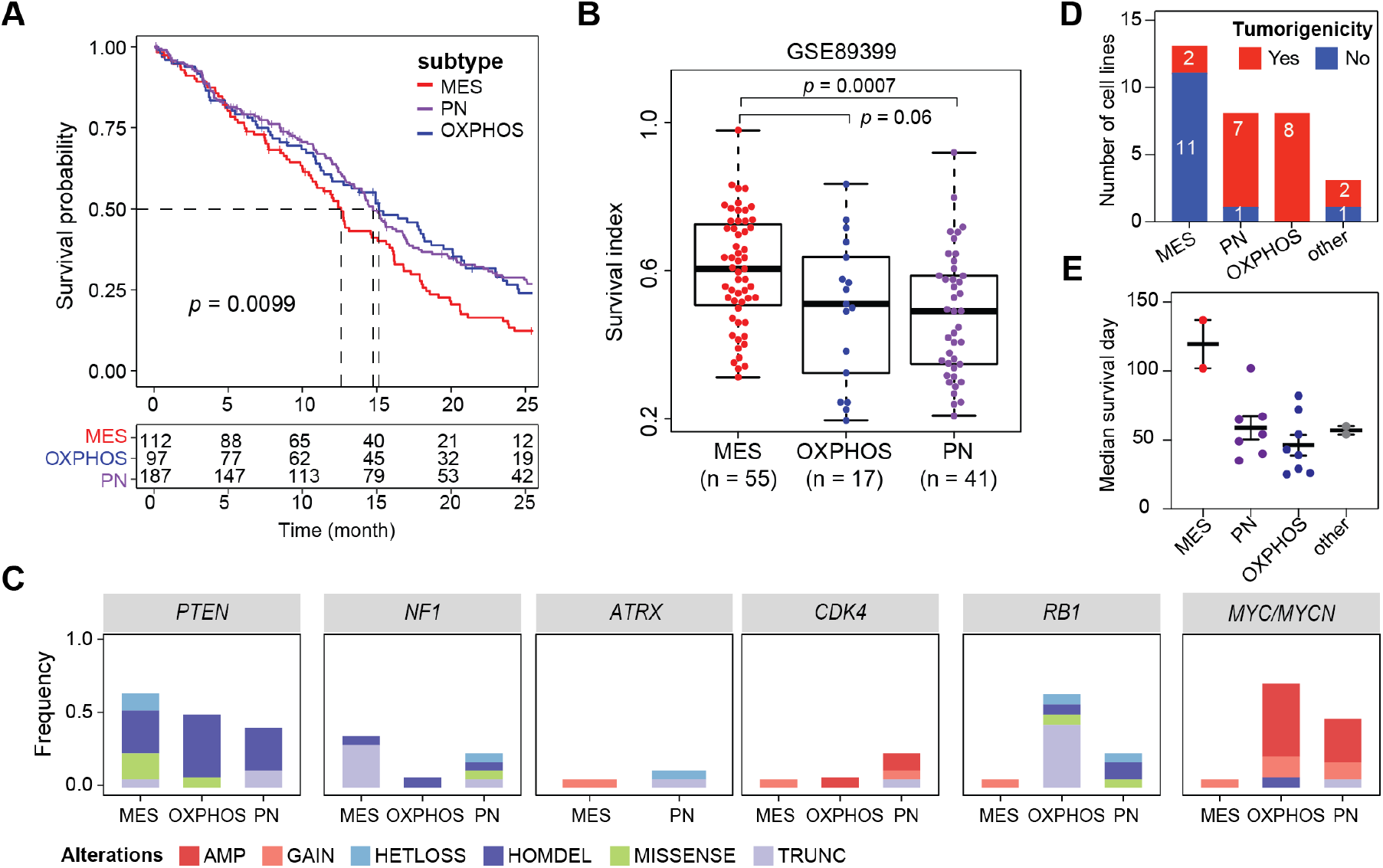
Clinical, genomic, and tumorigenic comparisons of defined subtypes, see also Figure S3. **(A)** Kaplan-Meier survival curves for GBM patients in the TCGA cohort with different subtypes. *P* value was calculated by log-rank test. **(B)** Radiotherapy response in defined subtypes. The y-axis represents the cell survival percentage upon radiation. *P* values were calculated by two-sided Wilcoxon rank sum exact test. **(C)** Genomic alteration frequency of indicated genes in PDGCs of different subtypes. Bars are colored by alteration types. AMP: high-level amplification, GAIN: low-level gain, HETLOSS: heterozygous deletion, HOMDEL: homozygous deletion, MISSENSE: missense mutation, TRUNC: truncation mutation. **(D)** Stacked bar plots showing the tumorigenicity of PDGCs of different subtypes. For each cell line, 5 × 10^5^ cells were injected into the brains of nude mice. **(E)** Scatter plots showing the median survival time of nude mice after injection of different subtype PDGCs. Lines were presented as mean ± SEM.

To evaluate the apparent radiotherapy sensitivity amongst the three subtypes we defined, we used the dataset from Segerman *et al*. (2016) study ^23^, wherein they established a library of primary glioma cell clones and investigated their gene expression as well as sensitivity to radiation therapy. After assigning the subtype identity to glioma cell clones, we found that the MES subtype cell clones were apparently more resistant to radiotherapy (median survival index: 0.61), while the PN subtype cell clones were relatively sensitive (median survival index: 0.49) (**Fig. 3B**).

To identify the genomic alterations in our subtypes, we detected copy number variations (CNVs) and mutations by analyzing WGS data of PDGCs (**Fig. 3C and Fig. S3C**). This analysis revealed that *PTEN* (MES: 64.7%, PN: 41.2%, OXPHOS: 50%) and *NF1* (MES: 35.3%, PN: 23.5%, OXPHOS: 7.1%) alterations were enriched in the MES subtype, *ATRX* (MES: 5.9%, PN: 11.8%) and *CDK4* (MES: 5.9%, PN: 23.5%, OXPHOS: 7.1%) alterations were enriched in the PN subtype, while *RB1* (MES: 5.9%, PN: 23.5%, OXPHOS: 64.3%) and *MYC/MYCN* (MES: 5.9%, PN: 47.1%, OXPHOS: 71.4%) alterations were enriched in the PN and OXPHOS subtypes (**Fig. 3C**). When checking the CNVs and somatic mutations of GBM patients from TCGA, we found a similar trend of genomic alteration distribution among our three subtypes (**Fig. S3C**).

To assess the tumorigenic ability of different subtypes of PDGCs, we transplanted 5 × 10^5^ PDGCs into the brains of immunodeficient mice. Most of the PN (7/8) and OXPHOS (8/8) subtype PDGCs were able to form intracranial tumors in nude mice (**Fig. 3D**). Despite GBM patients of MES subtype had the shortest overall survival compared to those of PN and OXPHOS subtypes (**Fig. 3A**), PDGCs of MES subtype had the lowest tumorigenicity (MES: 15.4%, PN: 87.5%, OXPHOS: 100%). Specifically, of the 13 MES PDGCs tested, only a fraction of mice transplanted with two cell lines (G709 and G98) formed intracranial tumors within 6 months after inoculation (**Fig. 3D and Table S5**). Furthermore, tumor-bearing mice transplanted OXPHOS subtype PDGCs tended to have shorter median survival than those transplanted PN subtype PDGCs (PN: 54 days, OXPHOS: 41 days) (**Fig. 3E**). Collectively, these findings revealed differences among the three subtypes in terms of apparent radiotherapy sensitivity, overall survival, genomic alterations, and tumorigenic ability.

### PDGCs with different subtypes have distinct drug responses

To explore the drug response of PDGCs of different subtypes, we initially evaluated the effects of 1,466 FDA-approved small molecules on the cell viability of three PDGCs (G98, G118, and G709). This screen yielded 214 small molecules that had at least a 75% inhibitory effect on at least one PDGC at 10 μM concentration (**Fig. 4A**). We subsequently measured the cell viability of 45 PDGCs treated with these 214 compounds (with a dose of 5 μM) (**Fig. 4A and Table S6**). We defined normalized cell viability (i.e., 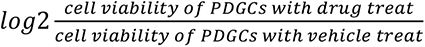) to measure the inhibitory effect of a drug to PDGCs. The lower the normalized cell viability, the stronger the inhibitory effects of the drug on the cells (**Table S7**).

**Figure 4.**
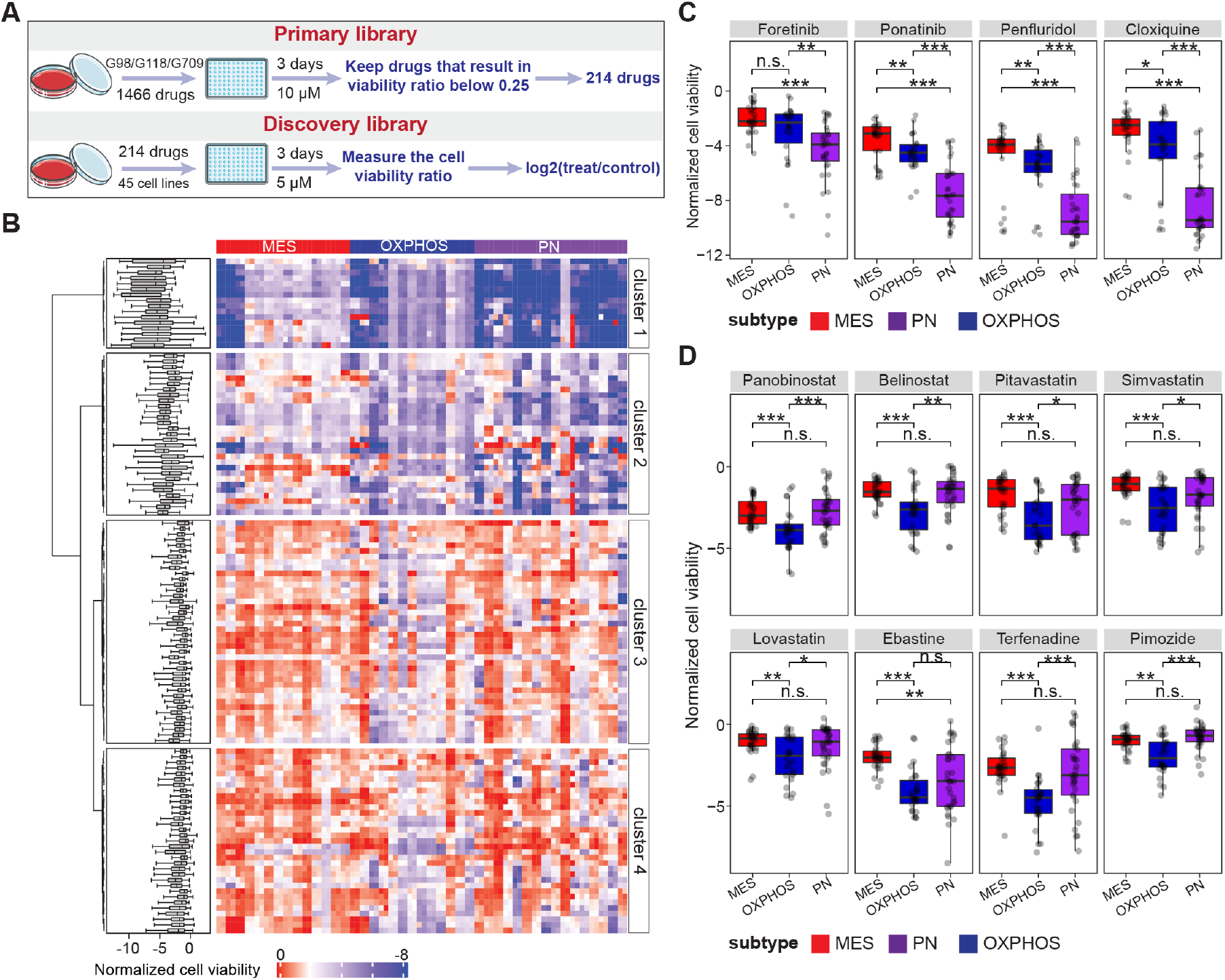
Drug responses of PDGCs of different subtypes, see also Figure S4. **(A)** Schematic diagram displaying the process of drug screening. **(B)** Heatmap showing the drug response of PDGCs of different subtypes. Rows represent drugs and columns represent cell lines, colored by the cell viability upon the treatment of corresponding drugs. Boxplots on the left side showed the response of cell lines to specific drug treatments. **(C-D)** Normalized cell viability of PDGCs of different subtypes upon treatment of the indicated drugs. Representative drugs that effectively inhibited the growth of PN and OXPHOS subtype PDGCs were listed in (**C**) and (**D**), respectively. *P* values were calculated by two-sided Wilcoxon rank sum exact test.

Cell viability analysis showed that the response of PDGCs to drugs varied, with the median normalized cell viability ranging from 0.033 to –9.76 (**Fig. S4**). We found that 96 of the drugs exerted little inhibitory effect on any PDGCs (normalized cell viability: mean > –1 or standard deviation < 0.5), so we retained 118 drugs for downstream analyses (**Fig. S4**).

To capture the drug response patterns across PDGC subtypes, we performed unsupervised clustering analysis on the cell viability profiles (**Fig. 4B**). Drugs in cluster 1 could inhibit cell growth across three PDGC subtypes, especially the PN subtype (**Fig. 4B**), such as anti-infection drug cloxiquine ^24^ (mean normalized cell viability: MES, –2.96; PN, –8.38; OXPHOS, –4.60), dopamine receptor blocker penfluridol ^25^ (mean normalized cell viability: MES, –4.97; PN, –8.85; OXPHOS, –5.78), and tyrosine kinase inhibitor (TKI) ponatinib ^26^ (mean normalized cell viability: MES, –3.59; PN, –7.45; OXPHOS, –4.54) (**Fig. 4C**). In addition, PDGCs with PN subtype were sensitive to drugs such as TKI (foretinib ^27^ and entrectinib ^28^), anti-infection drugs (cetrimonium bromide ^29^ and chlorhexidine 2HCl ^30, 31^), and antihistamine terfenadine ^32^ (**Fig. 4C and Table S8**). PDGCs with OXPHOS subtype were sensitive to drugs including mitochondrial activity inhibitors (Terfenadine ^33^ and Ebastine ^34^), histone deacetylase (HDAC) inhibitors ^35^ (panobinostat, belinostat, pracinostat, and abexinostat), and HMG-CoA reductase inhibitors ^36^ (pitavastatin, simvastatin, and lovastatin) (**Fig. 4D and Table S8**). These findings revealed that PDGCs of different subtypes showed differences in drug responses.

### PDGCs of OXPHOS subtype are sensitive to HMG-CoA reductase inhibitor lovastatin

Our drug screening showed that OXPHOS subtype PDGCs tended to be more sensitive to HMG-CoA reductase inhibitors than PN or MES subtype PDGCs (**Fig. 4D**). Upon treatment with lovastatin, OXPHOS subtype PDGCs had the lowest IC_50_ value among the subtypes (mean IC_50_: MES, 12.83 μM; PN, 1.98 μM; OXPHOS, 1.09 μM) (**Fig. S5A**). An Annexin V/PI assay of OXPHOS subtype PDGC BIN423 showed that treatment with 1 μM lovastatin led to a significant increase in the number of apoptotic cells (**Fig. S5B**). Statins block the synthesis of cholesterol by inhibiting HMG-CoA reductase (**Fig S5C**). We found that supplementing the lovastatin-treated BIN423 cells with cholesterol (or its precursor lanosterol) rescued the inhibitory effect of lovastatin (**Fig. 5A, B, and S5D**). These findings support that lovastatin inhibited the growth of OXPHOS subtype PDGCs by reducing the level of cholesterol.

**Figure 5.**
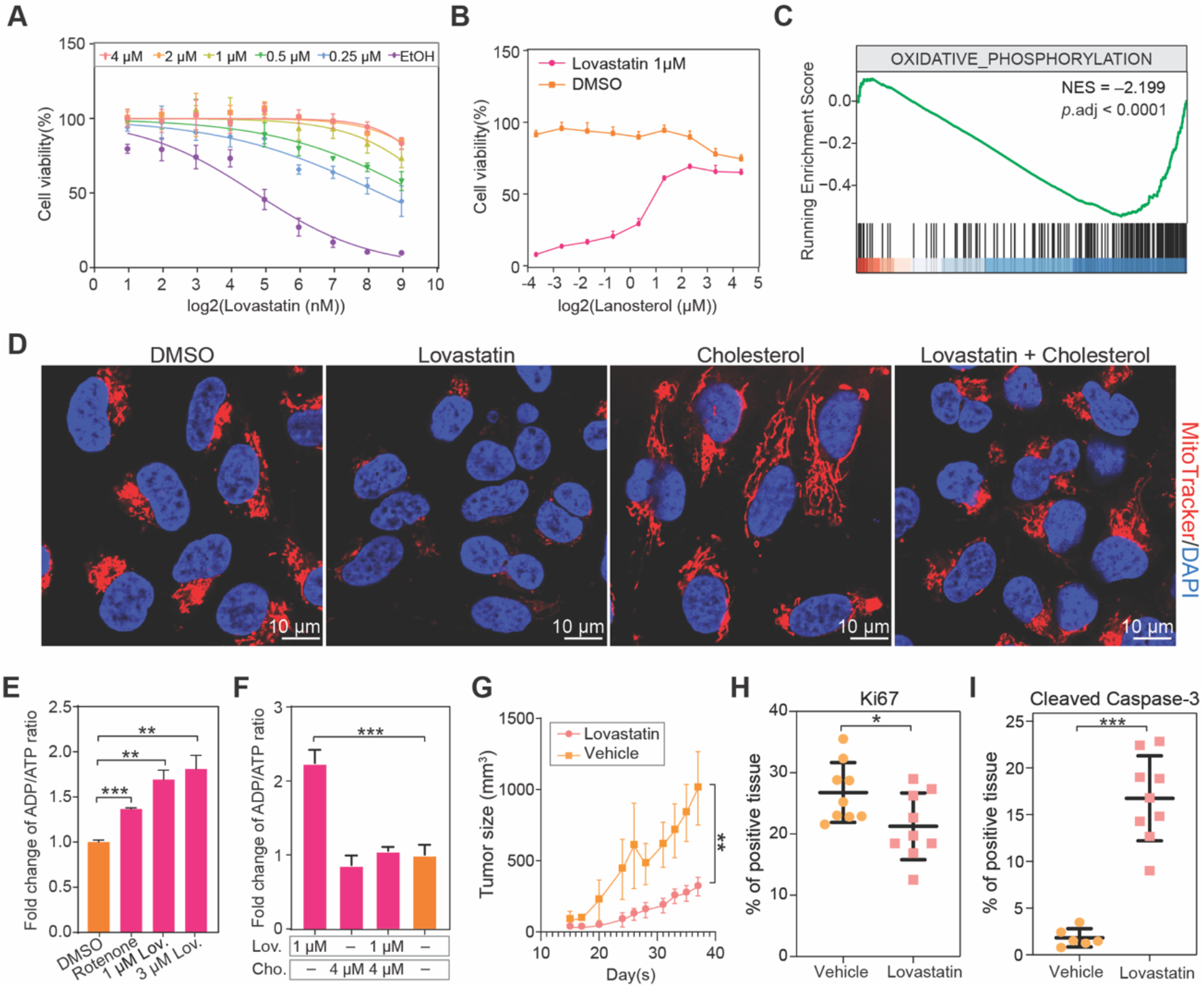
Lovastatin inhibits the growth of OXPHOS subtype PDGCs by attenuating mitochondrial respiration, see also Figure S5. **(A)** Cell viability of lovastatin-treated BNI423 supplemented with different concentrations of cholesterol (*n* =3). EtOH: ethyl alcohol. **(B)** Cell viability of lovastatin-treated BNI423 supplemented with different concentrations of lanosterol (*n* =3). **(C)** GSEA plot of OXPHOS subtype PDGCs treated with lovastatin compared to PDGCs without lovastatin treatment. GSEA is performed by R package clusterProfiler. **(D)** Representative MitoTracker staining of BNI423 under different treatments for 16 hours. Mitochondria were stained by MitoTracker (red), and the nucleus was stained by DAPI (blue). Scale bars, 10 μm. **(E)** Relative cellular ADP/ATP ratio of BNI423 under different treatments (*n* =3). Mitochondrial complex I inhibitor rotenone was used as a positive control. *P* values were calculated by two-sided Student’s t-test. Lov.: Lovastatin. **(F)** Relative cellular ADP/ATP ratio of lovastatin-treated BNI423 with or without cholesterol supplement (*n* =3). *P* values were calculated by two-sided Student’s t-test. Lov.: Lovastatin, Cho.: Cholesterol. **(G)** Tumor size of mice subcutaneously inoculated BNI17 (*n* = 6) treated with lovastatin or vehicle. *P* values were calculated by two-sided Student’s t-test. **(H-I)** Quantification of Ki67- (**H**) and cleaved Caspase-3-stained (**I**) brain tumor tissues. BNI423 were transplanted into the brains of nude mice. After five weeks, lovastatin was administered by brain infusion (42 μg per day per mouse) for three days, and the whole brain was excised for histopathological analysis. *P* values were calculated by two-sided Student’s t-test.

To investigate the downstream mechanism of lovastatin, we performed RNA-seq analysis of OXPHOS subtype PDGCs (BNI274, BNI423, and BNI7_11) with or without 1 μM lovastatin treatment, and a gene set enrichment analysis indicated that the oxidative phosphorylation pathway activity was significantly inhibited upon 18 h of lovastatin treatment (NES = –2.199, *p* < 0.0001) (**Fig. 5C**). This observation prompted us to check whether lovastatin treatment to OXPHOS subtype PDGCs would influence the number of mitochondria. We visualized mitochondria using MitoTracker and observed that the number of mitochondria was significantly reduced in BNI423 treated with lovastatin compared to the vehicle control group culture and this defect can be rescued by cholesterol (**Fig. 5D**).

Consistent with the RNA-seq analysis, we also found that treatment of BNI423 with lovastatin (Student’s t-test, *p* = 0.003) or with the mitochondrial complex I inhibitor rotenone ^37^ (Student’s t-test, *p* = 0.0002) resulted in significant increases in the ADP/ATP ratio compared to their respective vehicle controls (**Fig. 5E**). And again, the reduced ATP production phenotype upon lovastatin treatment was rescued by cholesterol supplementation (Student’s t-test, *p* = 0.0007) (**Fig. 5F**). Thus, lovastatin treatment of an OXPHOS subtype PDGC culture decreased the cholesterol level and thereby reduced the mitochondria number and reduced oxidative phosphorylation pathway activity. Daily oral gavage of lovastatin significantly reduced the tumor size of subcutaneously transplanted tumors (OXPHOS subtype PDGC BNI17) (**Fig. S5E and Fig. 5G**), but this regime was unable to prohibit the growth of intracranial tumors (**Fig. S5F**). Intracranial infusion of lovastatin through osmotic pumps (42 μg/day/mouse) significantly reduced mitotic index (**Fig. 5H and Fig. S5G**) (Student’s t-test, *p* = 0.038) and increased cleaved Caspase 3 staining intensity (**Fig. 5I and Fig. S5G**) (Student’s t-test, *p* < 0.0001) of transplanted brain tumors (OXPHOS subtype PDGC BNI423).

### Serum-free culture system expands cells with *MYC/MYCN* amplification

In the PDGCs we collected, 18 out of the 52 PDGCs (34.6%) harbored *MYC/MYCN* amplification and/or *MYC/MYCN* extrachromosomal DNA (ecDNA) amplification (**Table S1**). *MYC/MYCN* amplification is not a common event in GBM, only about 3.8% of GBM patients harbored *MYC/MYCN* amplification according to the data from the TCGA project ^38^. To explore whether MYC pathway activity is modulated by the cell culture system, we compared the transcriptomes of PDGCs and their matched tumor tissues and found the MYC pathway scores were significantly increased upon culture (**Fig. 6A**). We classified the 52 PDGCs into long (> 10 passages) and short groups according to their culture time (**Table S1**), and compared the frequency of *MYC/MYCN* amplification between the two groups. We found that the frequency of *MYC/MYCN* amplification events was significantly higher in long-cultured PDGCs than that in short-cultured PDGCs (48% vs. 22.22%, Fisher’s exact test, *p* = 0.048) (**Fig. 6B**). We observed that 36% of long-cultured PDGCs harbored *MYC/MYCN* ecDNA amplification, while this frequency in short-cultured PDGCs was only 11% (**Fig. 6C**). Besides, we analyzed 280 cell lines with WGS data from the CCLE project ^39^. *MYC* amplification occurred frequently in lung adenocarcinoma (LUAD), lung squamous cell cancer (LUSC), breast invasive carcinoma (BRCA), stomach adenocarcinoma (STAD), and ovarian cancer (OV) ^40^. Consistent with this notion, frequent *MYC* amplification was observed in their corresponding cell lines (percentage of cell lines having *MYC* amplification: lung, 20.41%; breast, 22.86%; stomach, 19.05%; ovary, 17.39%). No *MYC* amplification was detected in 15 CNS tumor cell lines (**Fig. S6A**), which were cultured under serum-containing conditions. These findings suggested that serum-free cell culture may promote *MYC/MYCN* amplification.

**Figure 6.**
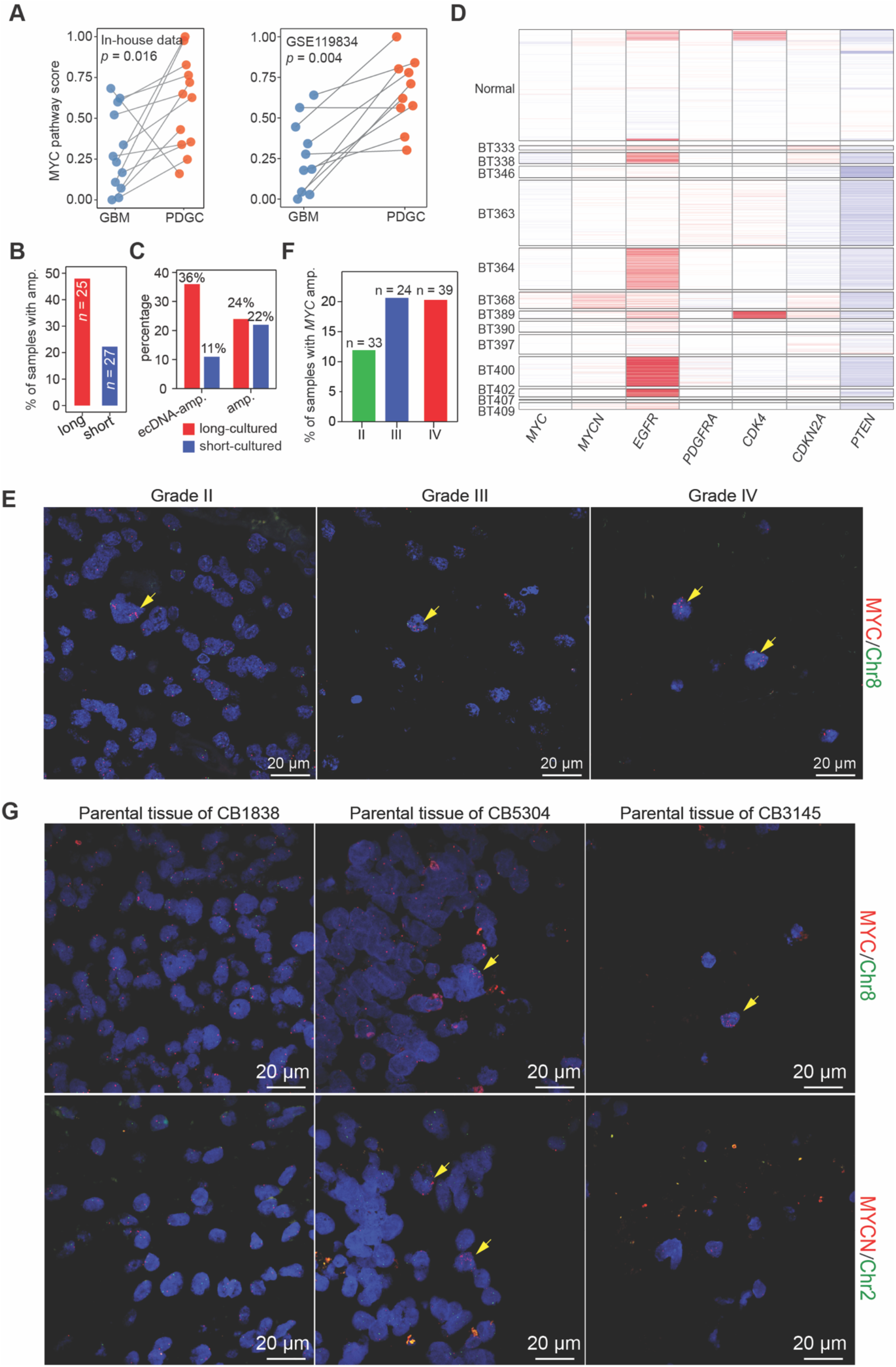
*MYC/MYCN* amplification in cultured PDGCs, see also Figure S6. **(A)** MYC pathway expression score in PDGCs and matched GBM tissues. *P* value was calculated by paired Wilcoxon rank sum exact test. **(B)** Percentage of samples with *MYC/MYCN* amplification in long- (*n* = 24) and short-cultured (*n* = 28) PDGCs. Cell lines that cultured more than 10 passages were determined as long-cultured, otherwise, the cell lines were determined as short-cultured. *P* value was calculated by one-sided Fisher’s exact test, *p* = 0.048. **(C)** Percentage of samples with *MYC/MYCN* DNA amplification or with *MYC/MYCN* ecDNA amplification. The red bar indicated long-cultured PDGCs, while the blue bar indicated short- cultured PDGCs. **(D)** Heatmap showing the CNV of indicated genes, colored by the CNV status (red for amplification and blue for deletion). Rows represent cells isolated from GBM patients and columns represent genomic locations. CNV levels were inferred by R package infercnv. **(E)** Representative FISH images showing rare MYC amplifying tumor cells in glioma tissues of grades II, III, and IV. The yellow arrow indicated cells with *MYC/MYCN* amplification. Scale bars, 20 μm. **(F)** Percentage of samples harboring more than three *MYC* amplification cells in glioma tissues of grade II (*n* = 33), III (*n* = 24), and IV (*n* = 39). **(G)** Representative FISH images showing the *MYC/MYCN* amplification in primary GBM tissues from patients CB1838, CB5304, and CB3145. The yellow arrow indicated cells with *MYC/MYCN* amplification. Scale bar, 20 μm.

As *MYC/MYN* amplification was also observed in low passage PDGCs, we speculated that a small number of *MYC/MYN*-amplified cells were presented in the original glioma tissues. To pursue this hypothesis, we initially analyzed scRNA-seq data of GBM tissues from Couturier *et al*. (2020) ^22^ and obtained 31,159 tumor cells after quality control. Inferred CNVs based on scRNA-seq data indicated that 38.8% of tumor cells harbored *EGFR* amplification and 50.8% of tumor cells harbored *PTEN* deletion. Notably, we found 1.2% and 5.4% of tumor cells harbored *MYC* and *MYCN* amplification, respectively (**Fig. 6C**). Upon analyzing scRNA-seq data of GBM tissues from Neftel *et al*. (2019) ^18^, we detected 4.6% of tumor cells harbored *MYC* amplification (**Fig. S6B**). We also examined additional 96 glioma tissue samples (33 grade II glioma tissues, 24 grade III glioma tissues, and 39 grade IV glioma tissues) by fluorescence in situ hybridization (FISH). We did detect cells with *MYC* amplification, and these cells tended to be distributed in clusters, suggesting that they may have originated from a single parental cell (**Fig. 6D, E**). Besides, high-grade glioma tissues had a higher probability of harboring cells with *MYC* amplification than low-grade glioma tissues (grade II, 12.12%; grade III, 20.83%; grade IV, 20.51%) (**Fig. 6E**). These findings collectively supported that there were rare cells with *MYC* amplification in glioma tissues.

If the observed *MYC/MYN* amplification in PDGCs was caused by the selective expansion of original tumor cells with *MYC/MYN* amplification, we would be able to locate these cells in the original tissues. To validate this, we performed FISH on matched tumor tissues of PDGCs. Our WGS analysis of PDGCs showed that CB5304 and CB3145 harbored *MYC* or *MYCN* amplification, while CB1838 did not harbor *MYC* or *MYCN* amplification (**Table S1**). FISH examination revealed that the parental tissues of CB5304 and CB3145 had cells with *MYC/MYCN* amplification, while the parental tissues of CB1838 did not have cells with *MYC/MYCN* amplification (**Fig. 6F**), suggesting that the *MYC/MYCN* amplification in PDGCs was not a *de novo* event. These findings collectively supported that the serum-free culture system expanded cells with *MYC/MYCN* amplification.

## Discussion

In this study, we integrated genomic, transcriptomic, and drug response profiles to characterize the glioma patient-derived cell lines. We classified the examined PDGCs into three subtypes: MES, PN, and OXPHOS. We found that the PN subtype PDGCs were sensitive to TKI, and OXPHOS subtype PDGCs were sensitive to HDAC inhibitors, oxidative phosphorylation inhibitors, and HMG-CoA reductase inhibitors.

We identified MES, PN, and OXPHOS subtypes based on the transcriptomes of PDGCs. The MES subtype was highly associated with MES^Wang^ subtype, suggesting the mesenchymal signature was stable in glioma cells. The PN subtype was enriched in PN^Wang^ and CLS^Wang^ subtypes. Studies in glioma mouse models demonstrated the subtype plasticity between the CLS^Wang^ and PN^Wang^ ^41^. The OXPHOS subtype was characterized by the high expression of genes involved in mitochondrial oxidative phosphorylation and resembled the mitochondrial (MTC) subtype defined by Garofano *et al*. ^42^; cells of this subtype exclusively relied on oxidative phosphorylation for energy production.

Our drug response analysis showed that the cell growth inhibition effect of multiple drugs was subtype-selective. For example, TKIs were especially effective against PN subtype PDGCs, which was consistent with a recent study that dasatinib, a multi-target TKI, conferred therapeutic benefit against oligodendrocyte lineage cells-derived GBMs ^43^. Oxidative phosphorylation inhibitors were particularly effective against OXPHOS subtype PDGCs. Compared to PN and OXPHOS subtype PDGCs, the MES subtype PDGCs were less sensitive to the screened drugs, which was consistent with the poor clinical prognosis of GBM patients with MES subtype.

We found that the cholesterol-lowering drug lovastatin prohibited the growth of OXPHOS subtype PDGCs by inhibiting cellular respiration *in vitro*. However, the anti-tumor effects of lovastatin *in vivo* were limited. In our study, oral administration of lovastatin was able to inhibit subcutaneous tumor growth, but brain infusion of lovastatin was ineffective for intracranial tumors. This is likely due to the fact that the brain is a cholesterol-rich organ, and the main source of cholesterol in the brain is *de novo* synthesis owing to the prevention of the blood-brain barrier ^44, 45^. Cholesterol is essential for fundamental neuronal physiology, and lovastatin treatment reduced the cholesterol level in the brain, which may influence the maintenance of normal brain function.

Compared to the traditional glioma cell lines cultured in a serum-containing medium, our PDGCs cultured in a serum-free medium more faithfully preserves the genomic alterations, intra- tumor heterogeneity, and transcriptional patterns of the parental GBM tissues. However, this culture system is not without flaws. We observed a high proportion of PDGCs with *MYC/MYCN* amplification, which is rare in GBM tissues. The presence of *MYC/MYCN* amplification had a strong correlation with the PDGCs culture system, which selectively expands the rare *MYC/MYCN-* amplified cells originating from the parental GBM tissues. Whether these *MYC/MYCN* amplified cells can still represent GBM needs to be seriously considered. For example, glioma cell line MES28 was originally isolated from tumor tissues of MES subtype ^46^ and widely used as an MES subtype cell line. However, after a long period of passaging, the MES28 has become OXPHOS subtype with *MYC* amplification.

We found that the majority of MES subtype PDGCs could hardly form transplantable tumors in immunodeficient mice. Thus, most *in vivo* studies involved glioma formation and treatment confined mainly to the PN and OXPHOS subtypes, while the MES subtype is relatively less studied and MES-specific compound has not been found by our drug screening. Multiple tumor cell subtypes co-existed in a single glioma tissue and only eliminating tumor cells of PN and OXPHOS subtypes could not effectively prevent tumor progression. This limitation may be one of the reasons for the low success rate of clinical translation from basic studies of glioma.

## Acknowledgments

This work was supported by the National Key R&D Program of China (2016YFA0503100, 2022YFA1103900) and the CAMS Innovation Fund for Medical Sciences (CIFMS) (2019-I2M-5- 015).

## Author contributions

M.W., R.Y., and J.C. conceptualized and designed this study. M.W. and J.C. wrote the manuscript. P.C., J.Z., and Y.L. constructed the PDGC lines. J.L. tested the tumorigenic ability of PDGCs. T.W. performed the initial drug screening. M.L. performed the drug screening experiments. R.Y. validated the effects of lovastatin *in vitro* and *in vivo*. M.W. performed bioinformatic analysis, drug response analysis, and generated figures. N.J., T.L., J.Z., G.L., Y.T., Z.G., X.W., A.W., W.C., M.G., G.C., Y.Y., and W.Z. provided the human glioma tissues or cell lines. Y.Y., W.Z., Q.W., and J.C. supervised the project and reviewed the manuscript. J.C. acquired financial support.

---

**Figure S1.**
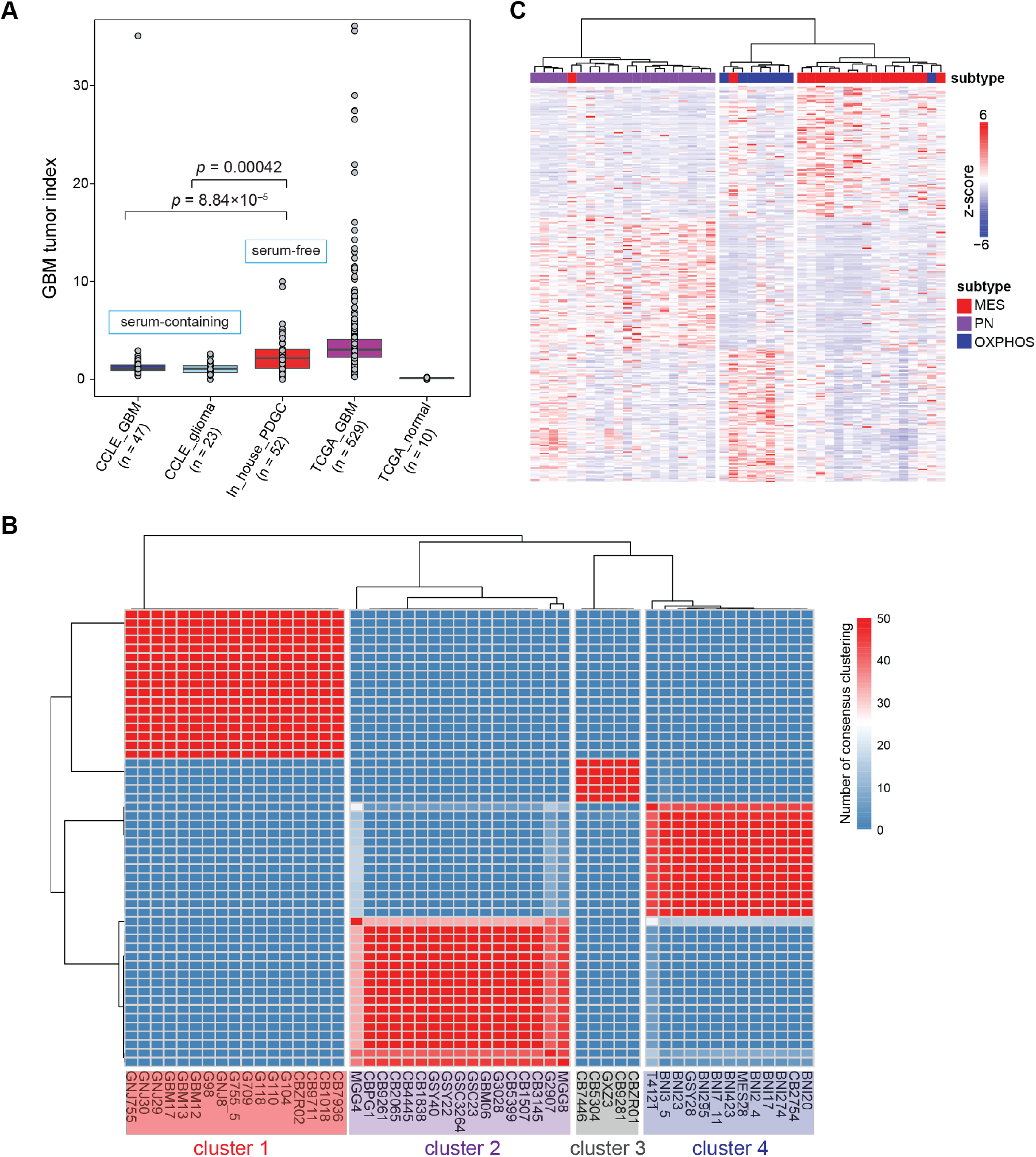
Definition of transcriptional subtypes for PDGCs, related to **Figure 1**. **(A)** Boxplot showing the GBM tumor index in cell lines and tissues. CCLE_GBM: GBM cell lines from CCLE (n = 47); CCLE_glioma: non-GBM glioma cell lines from CCLE (n = 23); In_house_PDGC: PDGCs cultured by serum-free medium (n = 52); TCGA_GBM: GBM tissues from TCGA (n = 529); TCGA_normal: non-tumor tissues from TCGA (n = 10). **(B)** Hierarchical clustering of 52 PDGCs. Rows represent genes, and columns represent cell lines. **(C)** Hierarchical clustering of 44 PDGCs in the GSE119834 dataset. Rows for the subtype-specific signatures and columns for the cell lines. Gene expression values were normalized by z-score.

**Figure S2.**
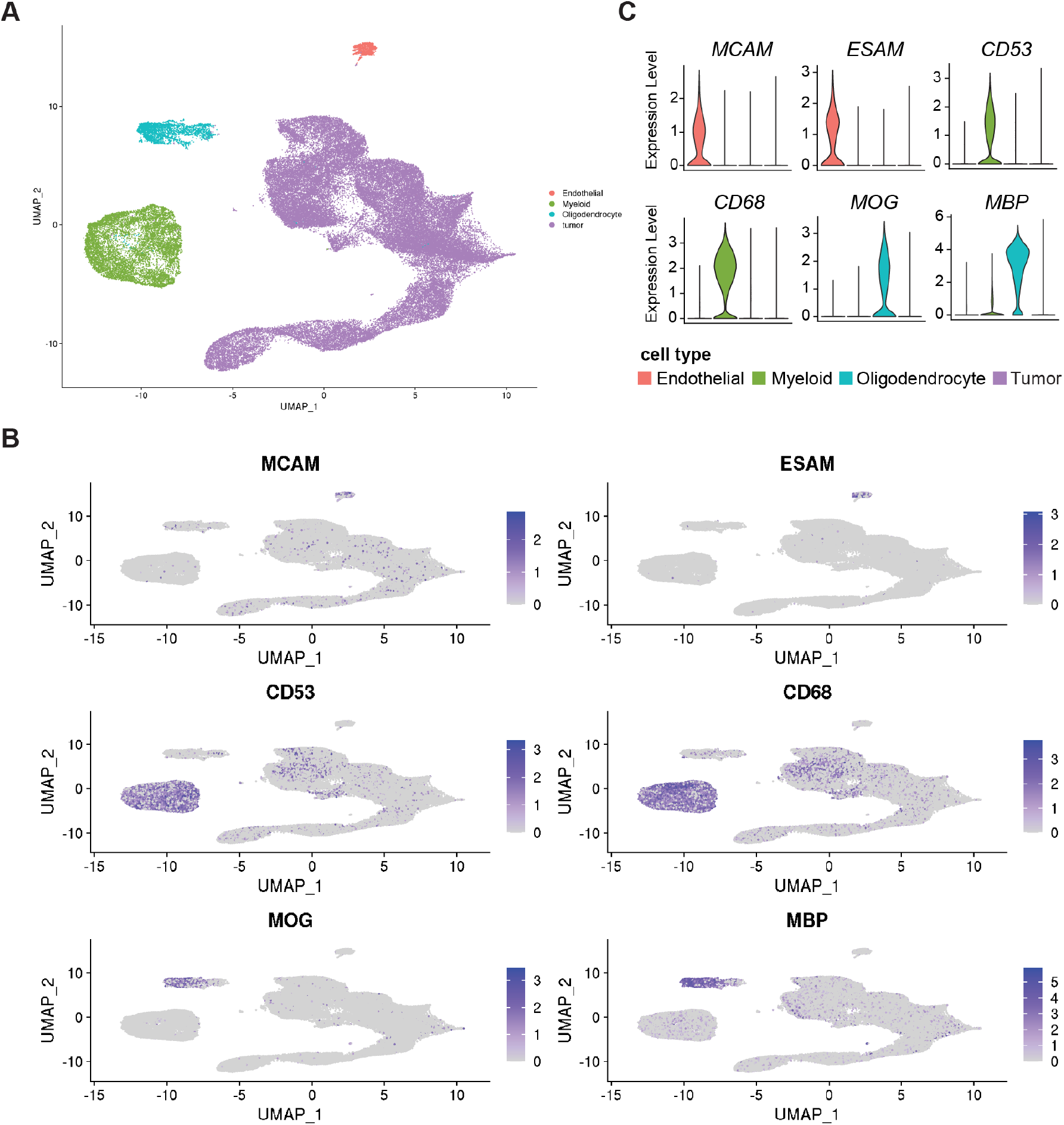
scRNA-seq data analysis for cell-type annotation, related to **Figure 2**. **(A)** UMAP plot of endothelial cells, myeloid cells, oligodendrocytes, and tumor cells. Each dot represents a single cell and is colored by cell type. **(B)** UMAP plots showing the expression of indicated markers. Cells were colored by the expression level. Endothelial cells: *MCAM* and *ESAM*; Myeloid cells: *CD53* and *CD68*; Oligodendrocytes: *MOG* and *MBP*. **(C)** Violin plots showing the expression of indicated markers in different cell types.

**Figure S3.**
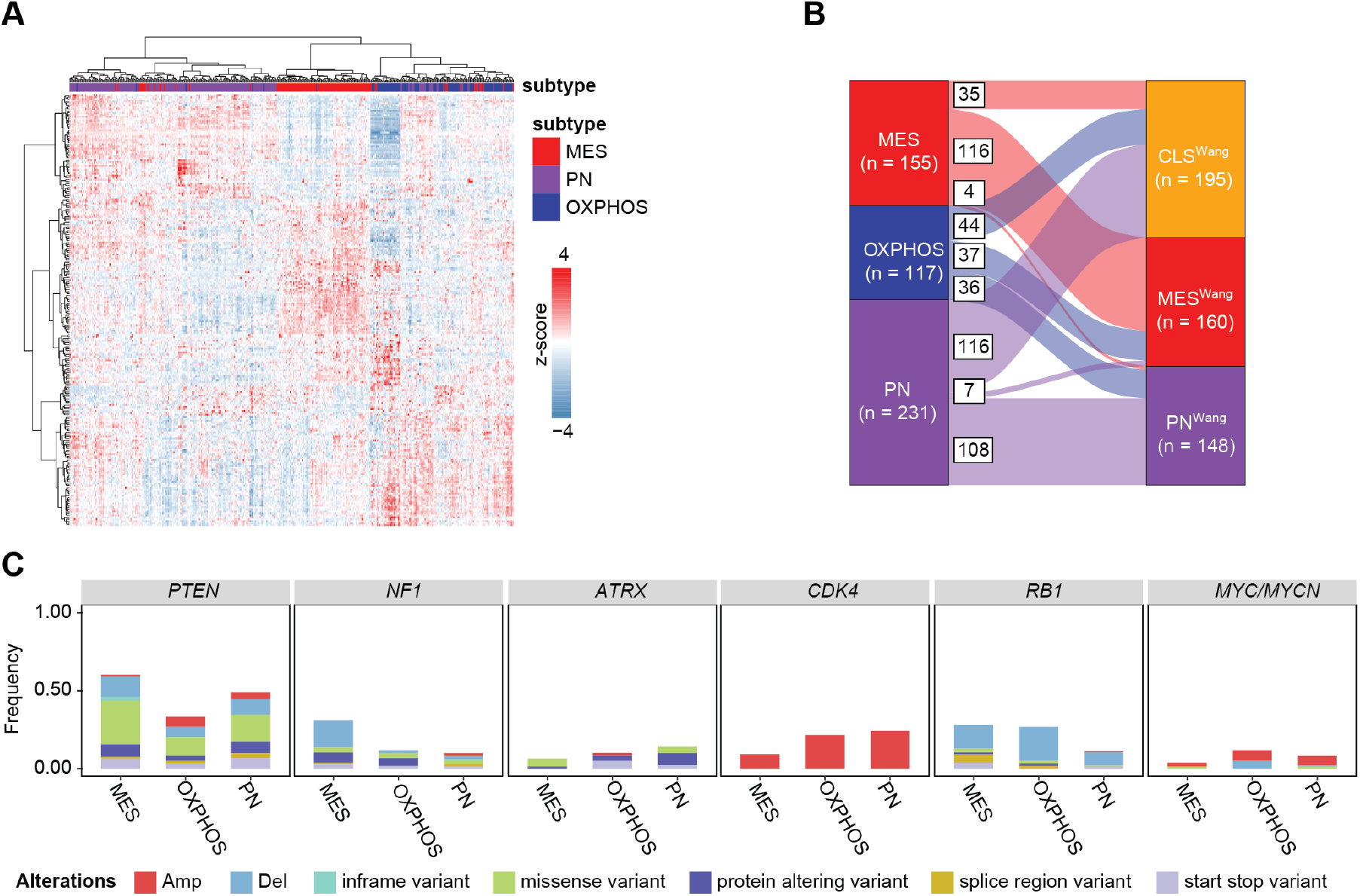
Validation of the transcriptomic and genomic characteristics of different subtypes in the TCGA cohort, related to **Figure 3**. **(A)** Hierarchical clustering of 403 GBM patients from the TCGA cohort. Rows represent the subtype-specific signatures and columns represent patients. Gene expression values were normalized by z-score. **(B)** Sankey plots showing the subtype assignment change flow. The left column is for our defined subtypes and the right column is for subtypes defined by Wang *et al*. (2017). **(C)** Frequencies of genomic alterations of indicated genes in different subtypes (TCGA-GBM cohort). Amp: amplification; Del: deletion; inframe variant: inframe insertion and deletion; protein altering variant: frameshift variant; splice region variant: splice donor variant, splice region variant, or splice acceptor variant; start stop variant: start lost, stop lost, or stop gained.

**Figure S4.**
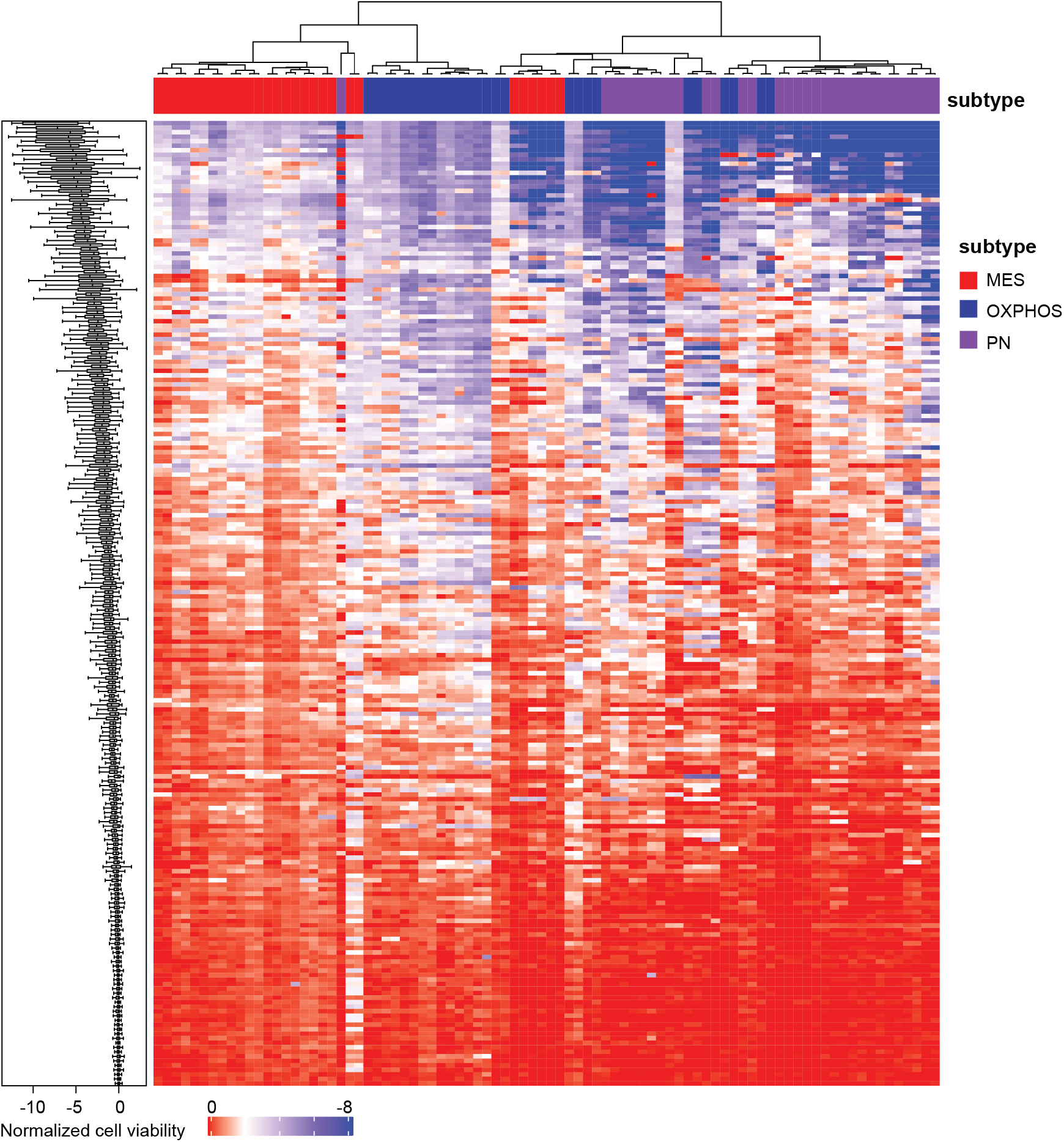
The drug response landscape of 45 PDGCs to 214 drugs, related to Figure 4. Heatmap showing the normalized cell viability of PDGCs upon drug treatment. Rows represent drugs and columns represent cell lines, colored by the cell viability upon the treatment of corresponding drugs. Boxplots on the left side showed the response of cell lines to specific drug treatments.

**Figure S5.**
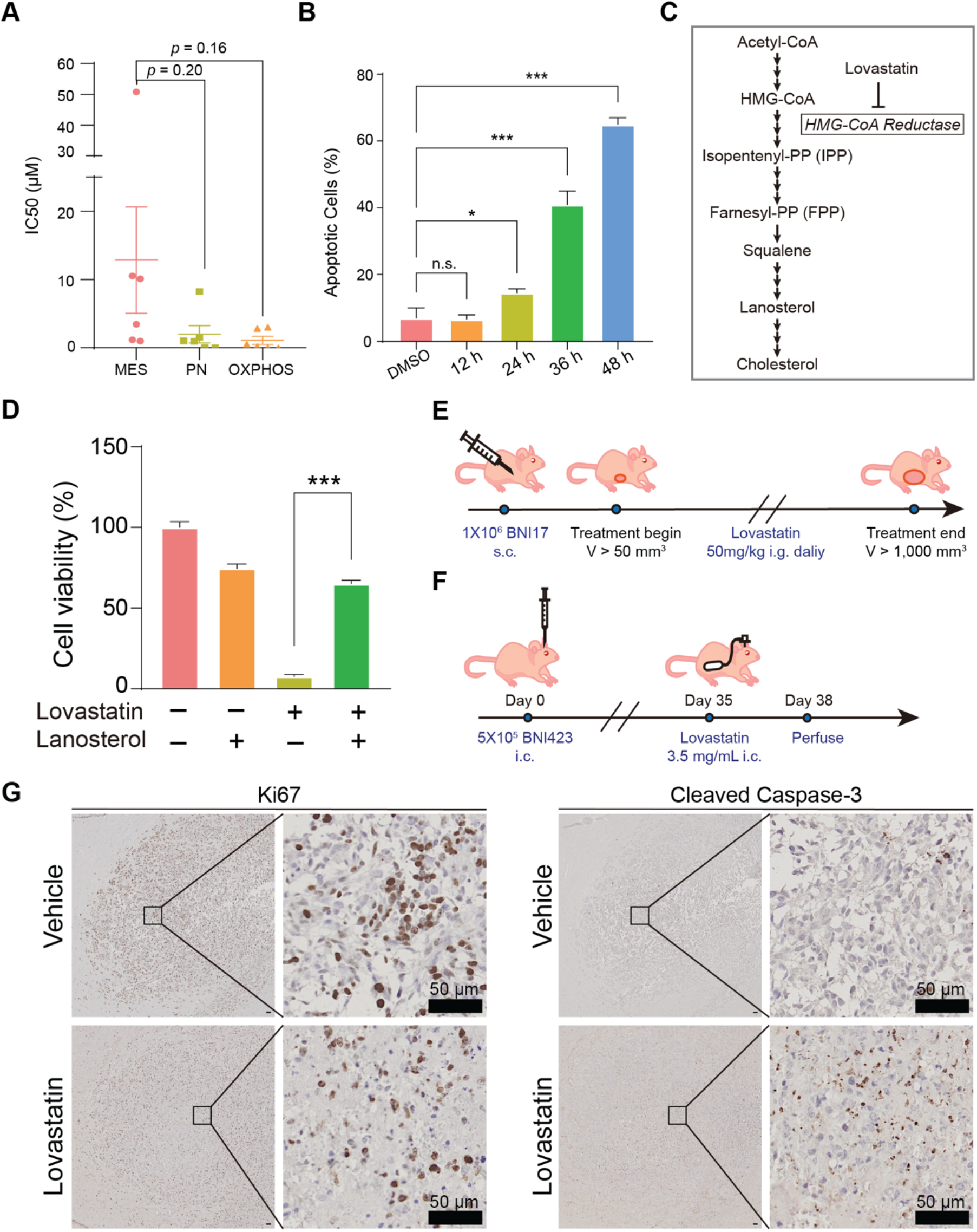
Lovastatin inhibits OXPHOS subtype PDGCs growth by attenuating mitochondrial respiration, related to **Figure 5**. **(A)** Lovastatin IC_50_ in PDGCs of different subtypes. MES (*n* = 6), PN (*n* = 6), OXPHOS (*n* = 6). *P* values were calculated by two-sided Student’s t-test. **(B)** Percentage of apoptotic cells in BNI423 after lovastatin treatment (*n* = 3). Apoptosis was evaluated by Annexin V staining. *P* values were calculated by two-sided Student’s t-test. **(C)** A simplified summary of the cholesterol synthesis pathway. **(D)** Cell viability of BNI423 treated with intermediate products in the cholesterol synthesis pathway. BNI423 cells were treated with lovastatin for three days and supplemented with cholesterol precursor products (i.e., lanosterol) (*n* = 3). *P* values were calculated by two-sided Student’s t-test. **(E)** Schematic diagram displaying the experiment design for subcutaneous xenograft. In brief, 1×10^6^ BNI17 cells were injected subcutaneously into mice. Lovastatin treatment begins when the tumor size of the control was between 50 mm^3^ and 1,000 mm^3^. **(F)** Schematic diagram displaying the experiment design for intracranial orthotopic xenograft. 5×10^5^ BNI423 cells were intracranially injected. After five weeks, the lovastatin treatment begins and continues for three days. **(G)** Representative IHC staining of Ki67 (left) and cleaved Caspase-3 (right) of intracranial tumor tissues. Scale bars, 50 μm.

**Figure S6.**
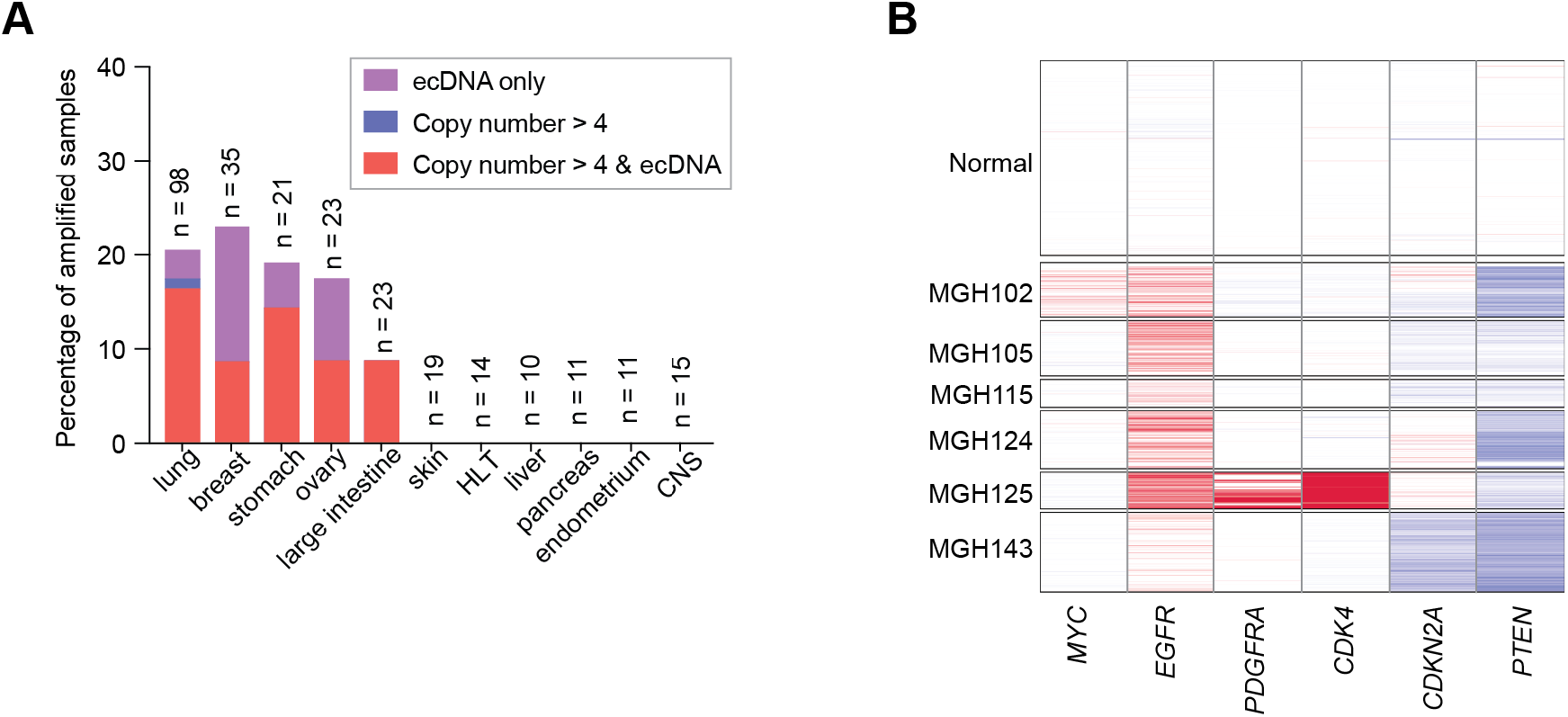
*MYC/MYCN* amplification in cultured PDGCs, related to **Figure 6**. **(A)** The percentage of *MYC* amplification in cell lines from the CCLE database. HLT, hematopoietic and lymphoid tissue. **(B)** Heatmap showing the CNV of indicated genes, colored by the CNV status (red for amplification and blue for deletion). Rows represent cells isolated from GBM patients and columns represent genomic locations. CNV levels were inferred by R package infercnv.

## Materials and Methods

### Collection and Processing of Patient-derived Glioma Cells (PDGCs)

Fresh glioma tissues were immediately placed in Hypo Thermosol FRS (Biolife Solutions, 101104) after surgery resection. The tissues need to be processed as quickly as possible because the longer the time between surgical removal and tissue processing, the more difficult it is for the PDGCs to survive. The tissues were transferred to a sterile tube and minced with dissection scissors in a biosafety cabinet. Accutase cell detachment solution (BioLegend, 423201) was added to the tube to resuspend the tissue pieces. To adequately disassociate tissues into single-cell suspension, the tube with the tissue pieces was incubated in a 37℃-water wash for 5∼10 min. Subsequently, the suspension in the tube was washed with cold PBS through a 40 µm cell strainer (Falcon, 352340) to remove un-disassociated tissue pieces. The cell precipitation was obtained by centrifuge at 500g for 5 min at 4℃. The supernatant was removed and red blood cell lysis buffer (Beyotime, C3702) was added with gentle pipetting at room temperature for 3∼5 min. Then centrifuged to remove the red supernatant. Washed the cells twice with cold PBS. The cells were resuspended in a serum-free culture medium and seeded into cell culture dishes.

### Cell Culture

The PDGCs were seeded into cell culture dishes (NEST, 705001), which were coated by Matrigel (CORNING, 354248), and cultured in a PDGCs serum-free medium containing DMEM/F12 (Gibco, C11330500BT), 1% penicillin/streptomycin (Solarbio, P8420/S8290), 1X B-27 without vitamin A supplement (Gibco, 12587010), 1X N-2 supplement (Gibco, 17502048), 0.5% glutaMAX-1 (Gibco, 35050-061), 5 mM HEPES (Beyotime, ST090), 600 ug/mL Glucose (Sangon Biotech, A501991-0500), 50 µM 2-mercaptoethanol (SIGMA, M3148-100ML), 20 ng/mL EGF (Novoprotein, C029-500 μg), and 20 ng/mL bFGF (ORIGENE, TP750002) and placed within a 37℃, 5% CO_2_, 5% O_2_, and 90% humidity sterile incubator. Usually, half of the medium was replaced every two days. When fully confluent, the PDGCs were split 1:2∼1:3 radios by Accutase cell detachment solution. In the early stage, PDGCs were frozen in each passage to preserve the original cell samples and expanded for further studies.

### DNA and RNA Extraction

For tissue samples from glioma patients, fresh-frozen tissue RNA was extracted by FastPure Cell/Tissue Total RNA Isolation Kit (Vazyme, RC112-01) according to the manufacturer’s instructions. For the PDGCs sample, 5×10^6^∼1×10^7^ cells were collected per cell line for DNA and RNA extraction. DNA of the PDGCs was extracted by TIANamp Genomic DNA Kit (TIANGEN, DP304), and RNA was extracted using standard TRIzol RNA extraction (Invitrogen, 15596-026). All nuclear samples were quantified using the NanoDrop instrument.

### Small Molecule Drugs Library Screening Assay

The small molecule library (Selleck) included 1,466 small molecule drugs approved by the FDA with a stock concentration of 10 mM in DMSO or water. These compounds were added to individual 96-wells at a concentration of 10 μM to select which of them would inhibit cell viability. This round of screening used three PDGCs (G98, G709, and G118). Small molecules with cell viability inhibition (cell viability below 0.25) in at least one cell line were selected for the second round of screening. In the second screening round, 45 PDGCs were tested at a concentration of 5 μM. Cell viability was determined using Cell Titer Glo ® (Promega, G9243).

### Cell Viability Assay

1,500 cells per well were seeded in 96-well plates. After 4∼6 h of incubation in a 37℃ incubator, cells were treated with different small molecule compounds as described in figure legends for three days. Cell viability was measured with Cell Titer Glo ® (Promega, G9243) after the treatment. Small molecule compounds purchased from Selleck included Lovastatin (S2061) and cholesterol (S4154), and from Shanghai yuanye Bio-Technology included lanosterol (S27466).

### Immunofluorescence (IF)

For mitochondria staining, BNI423 cultured on glass coverslips was incubated with lovastatin with/without cholesterol for 16 h. Subsequently, cells were incubated with 200 nM MitoTracker Deep Red FM (Invitrogen, M22426) for 40 min at 37℃, then washed in preheating PBS. Samples were fixed in PBS 4% PFA at room temperature for 1∼5 min and then permeabilized with PBST (PBS with 0.2% Triton X-100) for 10 min. Finally, samples were mounted with an antifade mounting medium with DAPI (Beyotime, P0131). Zeiss LSM880 confocal microscope with Airyscan was used to acquire images on a 63× oil objective.

### ADP/ATP Ratio Assay

For the cellular ADP/ATP ratio of BNI423, 3,000 cells/well were seeded in 96-well plates. After overnight incubation in a 37℃ incubator, the small molecule compounds were added to cultured cells. ADP/ATP Ratio was performed by ADP/ATP Ratio Assay Kit (SIGMA, MAK135) according to the manufacturer’s instructions after treatment.

### Apoptosis Assay

For Annexin V staining, BNI423 cells were seeded in 6-well plates with a density of 20%∼30%. After overnight incubation, cells were treated with lovastatin or vehicle (DMSO) for 12, 24, 36, and 48 h. FITC Annexin V/Propidium Iodide (PI) staining was performed by FITC Annexin V Apoptosis Detection Kit with PI (BioLegend, 640914). Flow cytometry was performed on BD LSR Fortessa (BD Biosciences) instrument, and data were analyzed using FlowJo (version 10.7.1).

### *In vivo* Test of Lovastatin

Female BALB/c Nude mice (6∼8 weeks) were purchased from Vital River and housed in the laboratory animal resource center (LARC) in the Chinese institute for brain research (CIBR), Beijing. All animal protocols were reviewed and approved by the Institutional Animal Care and Use Committee at CIBR. For the subcutaneous PDGCs’ xenograft model, 1,000,000 BNI17 (in matrigel) were subcutaneously injected into the left flanks of nude mice. For drug treatment studies, lovastatin (50mg/kg) or vehicle (PBS) was administered to mice by intragastrical (i.g.) daily after the tumor volume reached about 50 mm^3^. Tumor volume (V) was calculated with the formula V= (length (L)× width (W)^2^) ×0.5, where length and width were measured with the vernier caliper daily. Animals were observed until the tumor volume exceeded the limits (1,000 mm^3^). For the intracranial orthotopic PDGCs’ xenograft model, 500,000 BNI423 (in 5 μL matrigel) was transplanted stereotactically into the left striatum of nude mice by Standard Stereotaxic Instruments (RWD). The coordinates of PDGCs injection were + 0.5 mm AP, – 1.9 mm ML, and – 3.6 mm DV with respect to the bregma, and PDGCs were injected with a microsyringe (Hamilton, 701). Five weeks after the PDGCs transplantation, lovastatin (3.5 mg/mL) or vehicle was injected into the tumor site through the implanted brain infusion cannula. The brain infusion cannula (RWD, BIC-5) was connected via a 1.5 cm-long tubing to a subcutaneous osmotic pump (RWD, 1001W) to deliver the lovastatin or vehicle (flow rate 0.5 μL/h). After a three-day treatment, mice were perfused and whole brains were excised for further analysis.

### Histology and Immunohistochemistry (IHC)

The tissue was fixed in cold 4% paraformaldehyde (PFA) (Sangon Biotech, A500684-0500) overnight. The following day, fixed tissues were dehydrated through graded ethanol and embedded in paraffin. Paraffin blocks were sectioned at 3 µm intervals using a paraffin microtome (SLEE, Mainz, Germany) for H&E/IHC. The slides were deparaffinized and used the standard H&E staining protocol for histology. For IHC, antigen retrieval was performed in sodium citrate buffer (pH 6.0) in the pressure cooker. After washing three times with ddH_2_O, the slides were incubated in 3% H_2_O_2_ (Aladdin, H112515) for 10 min at room temperature. Then washing with ddH_2_O again and followed by blocking 3% BSA (SIGMA, V900933-100G) for 1 h at room temperature. Primary antibodies used included Ki67 (ORIGENE, TA802544) and cleaved caspase-3 (Cell Signaling Technology, 9661S). After overnight incubation of primary antibodies at 4 ℃, the slides were washed 4∼5 times with Tris Buffered Saline with Tween 20 (TBST). Chromogenic detection was performed with a two-step HRP-Polymer detection kit (ZSGB-BIO, PV-8000-1) and DAB kit (ZSGB-BIO, ZLI-9018). Olympus VS120 and ZEISS Axio Observer7 were used to acquire images and HALO was used for histopathological analysis.

### Fluorescence in situ hybridization (FISH)

We performed FISH on glioma tissues according to the manufacture’s instructions. *MYC* (8q24) gene amplification probe (LBP, F.01006) was purchased from Guangzhou LBP Medicine Science and Technology Co., Ltd, with chromosome 8 was labelled by centromere-specific probe. *MYCN* (2p24) gene probe (LBP, F.01013) was used to label *MYCN*, with a control probe to label *LAF4*.

### Bulk RNA-seq Data Analysis

Sequencing adapters of raw reads were trimmed by fastp ^47^ (version 0.23.1), and the resulting clean reads were mapped to the hg38 reference genome with STAR ^48^ (version 2.7.3a). Quantification of gene expression level was conducted by featureCounts ^49^ (version 2.0.0). The final gene expression matrix was normalized by counts per million (CPM).

### Bulk WGS/WES Data Analysis

Clean reads were aligned to the hg38 reference genome using bwa mem algorithm in Sentieon (Sentieon Inc, San Jose, CA). Germline mutations were called following the DNAseq pipeline in Sentieon (https://support.sentieon.com/manual/DNAseq_usage/dnaseq/), and somatic mutations were called following the TNseq pipeline in Sentieon (https://support.sentieon.com/manual/TNseq_usage/tnseq/). Mutations were annotated by annovar ^50^ (version 2019Oct24). Copy number variations (CNV) called by both CNVnator ^51^ (version 0.4.1) and hmftools-PURPLE (version 2.51) were retained for further analysis. In addition, extrachromosomal DNA (ecDNA) was analyzed by AmpliconArchitect ^52^ (version 2019).

### Subtype Determination Based on Non-Negative Matrix Factorization

We utilized the non-negative matrix factorization (NMF) clustering method to identify different subtypes among 52 glioma cell lines. Briefly, genes were ranked by the median absolute deviation (MAD) of expression values. We sorted MAD values from high to low and selected the top-rank genes for NMF clustering. When 52 glioma cell lines were clustered into four subtypes with the top 2500 genes, we received the highest cophenetic scores (**Table S3**). To generate stable clustering results, we used the “leave one out” method. Concretely, we only used 51 samples for NMF clustering each time. For example, we left the first sample out and used the 2^nd^ to 52^nd^ samples for clustering, and the next time, we left the second sample out and used the remaining samples for clustering. Finally, we obtained 52 clustering results (**Fig. S1B**). Given the fact that samples within the same subtype tended to cluster together, we determined the clustering identity based on 52 results.

### Identification of Subtype-specific Signatures

Subtype-specified signatures were identified as previously described ^42^. For gene *i*, we divided the expression values into two groups: A (what we are interested in) and B (the remaining samples). We defined a normalized enrichment score, NES, to estimate the level that the expression of gene *i* in group A is greater than the expression of gene *i* in group B.

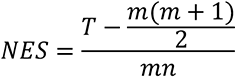

Where *m* is the number of samples in group A, *n* is the number of samples in group B, and *T* is the sum of ranks of gene *i* expression in group A. Next, we transformed *NES* into a fold change-based score: 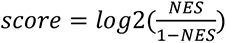). The higher the *score* is, the higher the expression of gene *i* in group A.

To identify the subtype-specific signatures, we calculated 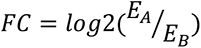, *p* values by Wilcoxon rank sum test, and adjusted *p* values by FDR, where *E_A_* is the average expression of gene *i* in group A, and *E_B_* is the average expression of gene *i* in group B. Signature of group A was determined by the following criteria: 1) *FC* > 1.5; 2) *p*. *adj* < 0.01; 3) top 100 ranked (from high to low) score.

### scRNA-seq Data Analysis

Couturier *et al*. data ^22^ was downloaded from https://github.com/mbourgey/scRNA_GBM. Basic analysis was performed by the R package Seurat ^53^ (version 4.0.3). Cells meeting the following criteria were retained for downstream analysis: 1) the number of detected genes was between 800 and 7500; 2) the number of unique molecular identifiers was between 1,500 and 50,000; 3) the percentage of mitochondrial counts was less than 10%; 4) the percentage of ribosomal counts was less than 30%. After quality control, cells from different patients were integrated by rliger ^54^ (version 1.0.0). Cells were annotated by canonical markers as indicated by the original article. *MCAM* and *ESAM* were expressed by endothelial cells, *CD53* and *CD68* were expressed by myeloid cells, *MOG* and *MBP* were expressed by oligodendrocytes. Transcriptome-based CNV inference was performed by R package infercnv ^55^ (version 1.6.0). Endothelial cells, myeloid cells, and oligodendrocytes were considered as references, with parameters “cutoff=0.1, denoise=TRUE, HMM=TRUE”. The resulting file run.final.infercnv_obj input to ComplexHeatmap ^56^ (version 2.6.2) for the visualization of CNV levels. Neftel *et al*. data ^18^ were downloaded from https://singlecell.broadinstitute.org/single_cell. Data were processed as above.

### GBM Tumor Index

TCGA-GBM gene expression data composed of 539 samples were downloaded from UCSC Xena^57^. Cancer cell line encyclopedia (CCLE) expression data were downloaded from DepMap (Public 22Q1) (https://depmap.org/portal/). We identified genes that were highly expressed in GBM tissues and defined these genes as GBM signatures. In the same way, we identified genes that were highly expressed in non-tumor brain tissues and defined these genes as non-tumor signatures. GBM signatures and non-tumor signatures were applied to CCLE cell lines and in-house PDGCs to calculate the respective signature score. The GBM tumor index was defined as the GBM signature score divided by the non-tumor signature score (Related to **Fig. S1A**).

### Calculation of Subtype Score

All the subtype signatures involved in this study were summarized in Table S3. Subtype score was calculated by R package ssGSEA ^58, 59^. Briefly, the ssGSEA score of each sample (for bulk RNA-seq) or cell (for scRNA-seq) was calculated and normalized to values between 0 and 1. In addition, we performed 1,000 permutations for each subtype signature and got the corresponding *p*-value. The subtype identity of each sample or cell was determined by the smallest p-value.

### Drug Response Data Analysis

We initially used a drug library composed of 1,466 FDA-approved small molecules to treat three PDGCs (G98, G118, and G709) respectively. After three days of exposure, we compared the cell viability between PDGCs with or without drug treatment. If the drug treatment (with a dose of 10 μM) inhibited cell viability (i.e., 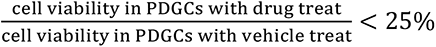), this drug was kept for further analysis. We obtained 214 small molecules that had the potential ability to inhibit glioma cell growth. Subsequently, we used these 214 drugs (with a dose of 5 μM) to treat 45 PDGCs for three days respectively, and measured the cell viability upon drug treatment. We defined normalized cell viability (i.e., 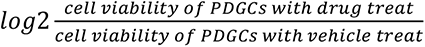) to measure the inhibitory effect of a drug to PDGCs. The lower the normalized cell viability, the stronger the inhibitory effects of the drug on the cells. Heatmap of the drug response data was generated by R package ComplexHeatmap ^56^ (version 2.6.2).

### Gene Set Enrichment Analysis

To detect the subtype-specific pathways, we download pathway information from MSigDB v7.3 (https://www.gsea-msigdb.org/gsea/msigdb/). Gene set enrichment analysis was performed by R package fgsea (version 1.16.0), with genes ranked by expression fold change. Visualization of enriched pathways was conducted by ggplot2 ^60^ (version 3.3.5).

### Survival Analysis

The survival data of TCGA patients were downloaded from UCSC Xena. The subtype of patients in the TCGA cohort was determined as above. Kaplan–Meier survival curves were generated by the R package survminer (version 0.4.9) to estimate the overall survival difference between subtypes, and *p* values were calculated using the log-rank test.

### Statistical Analysis

Detailed statistical information was listed in respective figure legends or methods. Asterisks are used to indicate the statistical significance (* *p* < 0.05, ** *p* < 0.01, *** *p* < 0.001); n.s. means statistically non-significant (*p* > 0.05).

### Data Availability

The RNA-seq, WES, and WGS data of PDGCs are available from China National Center for Bioinformation / Beijing Institute of Genomics, Chinese Academy of Sciences (https://ngdc.cncb.ac.cn) via accession numbers HRA003009.

